# SHOT-CCR: Biologically guided adversarial training for test-time adaptation in cellular morphology

**DOI:** 10.64898/2026.03.31.715531

**Authors:** William Dee, Aaron Wenteler, Srijit Seal, Otto Morris, Gregory Slabaugh

## Abstract

Pervasive batch effects are a common issue, especially in recent large-scale Cell Painting datasets, which have been produced to aid AI-enhanced drug discovery efforts. Technical differences arising from experiments carried out in different batches can cause models to fail to generalize to unseen batches, despite good predictive performance “within batch”. We propose a biologically grounded test-time adaptation framework, SHOT-CCR, which uses cell-invariant gradient reversal to decouple morphological signal from experimental confounders. Our approach performs 4.5% better than the current RxRx1 benchmark, classifying 1,139 classes of siRNA genetic perturbations with 91.6% accuracy. We deliver consistent results over four distinct cell types and two prominent Cell Painting datasets – RxRx1 and a subset of JUMP-CP. Across 484 classes of CRISPR perturbations in JUMP-CP our method improves accuracy by 15.7%.

## 1 Introduction

Over the past decade the availability of high-content screening (HCS) data has grown dramatically [30, 5, 6, 7]. High-throughput microscopy screening systems are now an important component within drug discovery, enabling biologists to image the responses of millions of cells to thousands of different chemical and genetic perturbations [15, 10]. This capability is relatively cheap when compared with mRNA or protein profiling and contributes distinct information regarding a cell’s phenotypic response [1, 8]. Computer vision pipelines have been leveraged for a variety of cellular perturbation tasks, including discerning molecular mechanism of action [8, 22, 24, 28], compound toxicity [27, 26], gene function and disease mechanisms [1].

Multiple studies have documented how technical artifacts and other confounding batch effects can arise even under the most stringent experimental settings [8, 22, 24, 26]. To compound this, larger datasets necessitate a greater number of separate experimental batches. These batch effects typically obscure biological cellular processes, causing models to fail to generalize to out-of-batch samples.

Sypetkowski et al. [30] showed that, in the large and diverse Cell Painting dataset RxRx1, adaptive batch normalization (AdaBN) can be an effective strategy to mitigate the impact of batch effects across experimental batches for certain cell types. Their model achieved a classification accuracy of 87.1% across 1,139 unique siRNA perturbations, with the model being able to generalize with 95.6% effectiveness to unseen batches. However, the approach underperformed for the U2OS cell type, which accounted for less than 10% of the training data, classifying those perturbations with 68.2% accuracy.

To improve these results, we extend domain adaptation techniques, originally developed for natural image analysis, to the problem of experimental batch variation in Cell Painting data. We incorporate recent advances such as entropy minimization, diversity regularization, and pseudo-labeling, and combine them with adversarial training designed to de-prioritize the cell count differences which commonly arise across experimental batches and cell types. Crucially our method, SHOT Cell Count Reversal (SHOT-CCR), does not aim to eliminate all batch information from the encoder embeddings. Instead, it selectively down-weights cell-count–driven batch differences while retaining residual batch-specific structure that remains predictive of the underlying perturbation. We focus on cell count in this work due to the fundamental biological nature of the feature and how integral it has proven to be in other Cell Painting research [5, 8, 28, 26]. Our main contributions are as follows:

1. **Biologically informed test-time adaptation**. We propose a framework that extends test-time adaptation techniques from computer vision to Cell Painting data, resulting in robust generalization under unseen batch and cell type shifts.
2. **Cell count adversarial training**. We introduce a novel biologically guided adversarial mechanism that reduces batch effects by discouraging the network from over-relying on cell count features—outperforming the general batch-effect gradient reversal of Sypetkowski et al. [30].
3. **Comprehensive evaluation across datasets and cell types**. We demonstrate consistent gains across two large-scale Cell Painting datasets (RxRx1 and JUMP-CP) and four distinct cell types, establishing a new benchmark for morphological batch correction.

## 2 Related Work

### Cell Painting batch effect correction

The reduction of batch effect impact is a central theme in most Cell Painting microscopy research [30, 5, 6, 7, 15, 8, 37]. Image embeddings and other downstream representations, such as CellProfiler image-based profiles [4], have been shown to group tightly together according to these effects, a phenomenon which makes classification tasks difficult. There is a growing desire to amalgamate microscopy datasets from either different sources (e.g. labs) or different time periods (to utilize legacy data). The belief is that more diverse data will result in robust models which will generalize more effectively to cell types, drug compounds and genetic perturbations which have not been extensively studied, facilitating more scalable and efficient drug/treatment discovery [5, 6, 7, 15]. To realize this vision there needs to be a universal approach to batch effect correction proven to be effective across a variety of Cell Painting datasets, and which shows the ability to recover meaningful biological signal from the images while reducing confounding effects.

### Test-time adaption (TTA)

TTA refers to methods that enable models to adapt to new, previously unseen data distributions during inference without requiring model re-training or access to the original training data [19]. There are several publicly available large-scale Cell Painting datasets [30, 5, 6, 7, 9], as well as released models which have been pre-trained on those datasets [30, 15, 28]. It is therefore becoming increasingly necessary to explore TTA methods which enable researchers to apply previously trained models to new experimental batch data with potentially different cell types present. The simplest TTA approaches either recalculate batch normalization statistics using target domain data, or update batch normalization parameters through entropy minimization [17, 35]. More advanced methods also add pseudo-class labelling or prototypes (weighted averages of test features) based on predictive class confidence either on the original test set images or augmented versions [18, 36, 14]. Model adaptation can take place either through a series of episodes (i.e., test batches), via a single-pass epoch or, as we have done in this work, throughout iterative, multi-epoch adaptation using self-supervised objectives.

### Gradient reversal

is a technique used for learning domain-invariant representations that retain relevant information for the goal task while minimizing the impact of one or many domain-related variables. To achieve this, model gradients flowing back from a domain-specific head are multiplied by a negative factor to encourage worse predictive performance for that variable [2, 11, 34]. Our work utilizes gradient reversal to selectively reduce, but not eliminate, the impact of cell count differences across multiple experiment batches in our data. We show there is a trade-off between eliminating the impact of a batch-related variable like cell count and classification accuracy. We demonstrate that adding gradient reversal learning parameters can improve model accuracy during training and improve subsequent TTA performance.

## 3 Datasets

### 3.1 RxRx1

RxRx1 was produced by Recursion using their high-throughput screening platform [30]. In each experimental well, cells are genetically perturbed with a fixed concentration of small interfering ribonucleic acid (siRNA), designed to “knockdown a single target gene via the RNA interference pathway, reducing the expression of the target gene” [30, 33].

The authors applied a proprietary implementation of the original Cell Painting protocol [3], producing a six-channel image. Each channel corresponds to a fluorescent dye which aims to represent the response of specific cellular components, including the nucleus, endoplasmic reticulum, actin, nucleoli, mitochondria and Golgi (see 1 example). The dataset was intended to serve as a baseline for future authors to compare batch correction methods against.

RxRx1 contains 51 experimental batches, split across four separate cell types—HUVEC (24 batches), RPE (11), HepG2 (11) and U2OS (five), where each batch experiment was conducted a minimum of one week after the last. Of these, U2OS was considered the most biologically distinct and therefore most difficult to learn to predict, especially in a limited data scenario [30]. We use the same batch-separated splits as Sypetkowski et al. [30], where 33 experimental batches were assigned to the training set (16 HUVEC, 7 RPE, 7 HepG2, 3 U2OS) and the remaining 18 were assigned to the test set (8 HUVEC, 4 RPE, 4 HepG2, 2 U2OS).

### 3.2 JUMP cpg0016 (JUMP-CP)

We filtered the JUMP-CP project dataset to only include perturbations where genes were knocked out using Clustered Regularly Interspaced Short Palindromic Repeats (CRISPR) guides [6, 7]. We then selected only genetic perturbations that were present in more than one batch, and where there were at least five replicates performed across five separate plates. This ensured there was adequate data for the model to learn from for each perturbation. For comparability with Sypetkowski et al. [30], we selected only genes which were present in the RxRx1 data, resulting in 484 unique genetic perturbations spread across five experimental batches (1). Only the U2OS cell type is used in JUMP-CP, and the original Cell Painting protocol produces five-channel images, rather than six [3]. Despite this, the stains aim to display functionally similar cellular components to the RxRx1 dataset.

Due to the smaller diversity of data when compared with RxRx1 (1), and the known difficulty in predicting CRISPR knockouts compared to siRNA knockdowns [16], we expect the models to perform more poorly on this data.

## 4 Methodology

### 4.1 Model architecture

An overview of our model architecture is shown in 2. The architecture is based on the DenseNet-161 backbone proposed in the original RxRx1 paper [30]. The backbone is adapted for Cell Painting images by altering the first convolutional layer to accept either five (JUMP-CP) or six-channel (RxRx1) images as input, before a Kaiming normal initialization [12] is used to initialize the additional channel weights. Global adaptive average pooling is applied to the backbone feature maps before passing through two fully connected layers, each utilizing batch normalization and the ReLU activation function.

The model is trained for 100 epochs, applying horizontal and vertical flips, and 90-degree rotation augmentations with a probability of 0.8. We applied CutMix [38] to all training images with a probability of 1.0, applying proportional mixing of auxiliary features (i.e., cell counts) based on the spatial mixing ratio. The images are self-standardized channel-wise, i.e., each image channel is standardized to have a mean of zero and unit variance. We used an effective batch size of 512 for training, equating to two Nvidia A100 GPUs, processing batches of 32 each with gradients accumulated for eight steps.

We also evaluated EfficientNetV2 [32] and ConvNeXt [21] as alternative backbones to benchmark the architectural performance against the established DenseNet161 approach. However, as this comparison is not the primary focus of the study, all main results, unless otherwise specified, are reported using DenseNet161 to maintain comparability and consistency with Sypetkowski et al. [30].

### 4.2 Cell-count gradient reversal (CCR)

In contrast to Sypetkowski et al. [30], we use a gradient reversal layer to adversarially train the model to de-prioritize cell count. A separate regression head, consisting of two linear layers, is used for this task and the gradient reversal layer feeds back the inverse of the loss to the feature extractor (Figure 2).

**Figure 1.**
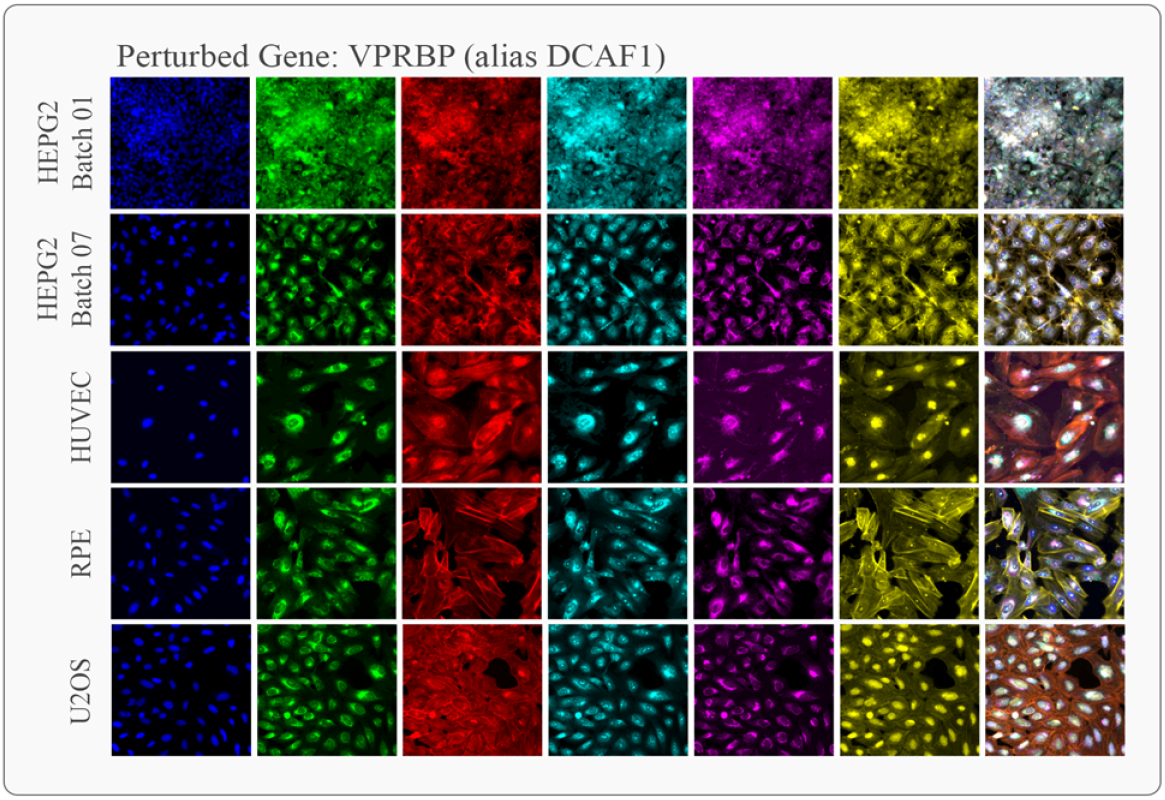
Six-channel faux-colored images from the RxRx1 dataset with a composite image shown on the right. Different stains reveal a range of cellular components across each channel (columns). Each set of images displays the same perturbation class (a knockdown of the VPRBP gene) but imaged in different batches and/or different cell types. The variety of cellular responses to the *same perturbation*, which can be seen by eye above, show how difficult it can be for a model to adapt to both biological *and* technical batch effects.

**Figure 2.**
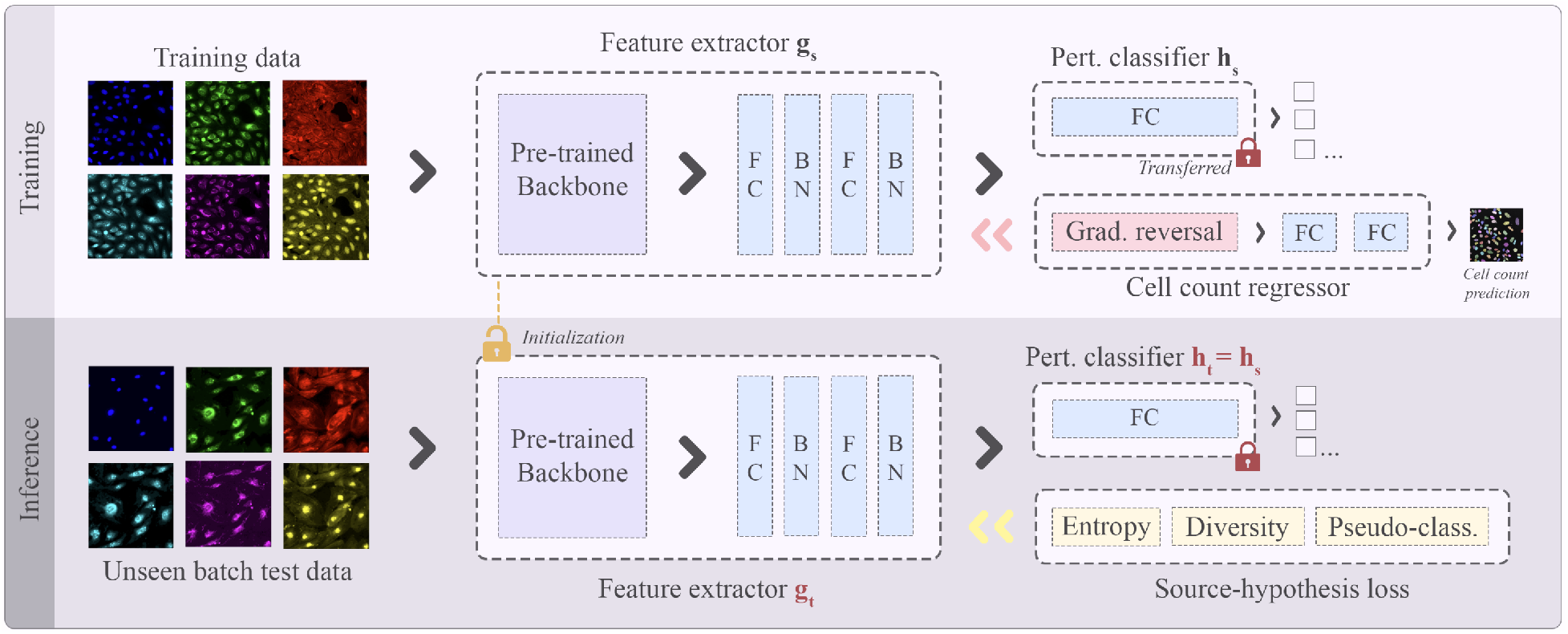
A diagram of our model architecture. The backbone network can be any pre-trained model which utilizes batch normalization layers, we found both DenseNet161 [13] and EfficientNetV2 [32] to be effective. Feature maps from the backbone are pooled using global averaging before being passed to two fully connected layers, each with batch normalization layers. During training the feature extractor (*g*_*s*_) is connected to a linear layer (the “perturbation classifier”, *h*_*s*_), which predicts the perturbation class, and a cell count regression head which uses gradient reversal to adversarially train the model feature maps to be less aligned to image cell count. At test-time the perturbation classifier is frozen while the feature extractor is trained further in an unsupervised manner using source-hypothesis (SHOT) loss [18]. Unless otherwise stated, all results in this paper utilize the DenseNet-161 backbone, for comparability with Sypetkowski et al. [30].

Since cell count is trivial for the model to learn to predict, we introduce separate learning rate and gradient reversal alpha (*α*) parameters to avoid both over-fitting as well as training the model to be completely agnostic to cell count – both of which appear detrimental to model performance (see Section 5).

The cell count in each image was obtained by feeding the nuclei-stained channel through CellposeSAM (cellpose v4.0.1) [23]. The cell count regression head predicts normalized cell counts of each image during training. We calculate normalized cell counts using the mean and standard deviation of cell counts in the training data. We use mean-squared error (MSE) loss to measure the degree of cell-count invariance reflected in the models’ embeddings during training. MSE is calculated by comparing the normalized model cell count predictions to the actual cell counts from Cellpose. During inference and test-time adaptation cell counts were not used to avoid any form of data leakage.

### 4.3 Test-time adaptation

#### Adaptive batch normalization (AdaBN)

AdaBN [17] utilizes batch normalization layers to standardize samples only using other samples produced in the same domain. In practice, this means that during training a sampler is used to ensure that each mini batch only contains data from a single domain (an experimental batch in our case) - i.e., batch effects normalization (BEN [20]). At inference time, the running statistics accumulated on the training set are discarded and instead, normalization is performed according to test mini batch statistics (which are also sampled from a single domain).

#### Test entropy minimization (Tent)

Tent [35] adapts a pretrained model solely at inference time rather than altering the model approach during training. The approach aims to minimize the Shannon entropy [29], i.e., uncertainty, of model predictions, gradually increasing model prediction confidence throughout adaptation. Unlike AdaBN, Tent adapts both the scale (*γ*) and shift (*β*) parameters of the model’s batch normalization layers during test-time through the backpropagation of entropy loss. In this manner, the method remains unsupervised since it relies only on model predictions, not labels, to update model parameters.

#### Source hypothesis transfer with information maximization (SHOT)

SHOT builds on the Tent approach but instead freezes the classifier module (the hypothesis) from the pretrained model and uses the feature extractor module(s) as initialization for target domain learning during test-time adaptation [18]. To adapt the feature module, Liang et al. [18] proposed a combination of entropy minimization loss, ℒ_ent_, diversity loss, ℒ_div_, and self-supervised pseudo-labeled classification loss, ℒ_pc_.

ℒ_ent_ and ℒ_div_ constitute the information maximization loss, which aims to make target outputs both “individually certain and globally diverse” [18]. The diversity term aims to ensure model predictions don’t become overly confident and narrow to focus on a few specific classes as entropy loss is reduced. Within a batch, if model confidence exceeds a threshold, beta (*β*), a pseudo-label is given to the sample based on that prediction and the classification cross entropy loss for that sample is calculated. This pseudo classification loss (ℒ_pc_) is then subtracted from the information maximization loss (ℒ_ent_ + ℒ_div_).

In our work we freeze the perturbation classifier linear layer of our model after initial training (see 2). The trained feature extractor is used for initialization at the beginning of test-time adaptation and then its weights are updated based on feedback from the unsupervised SHOT loss. For test-time adaptation we use a learning rate of 1e-5, the Adam optimizer and a confidence threshold *β* of 0.95. We train over multiple epochs, using an early stopping criterion of 15 epochs and select the model where SHOT loss reaches a minimum.

#### Other test-time adaptation methods

Additional test-time adaptation methods have proven suitable for domain adaptation in other fields within computer vision. However, we found that not all methods were suitable for Cell Painting data. Continual test-time domain adaptation (Cotta) uses a teacher-student model framework whereby the teacher is presented with multiple augmented views of an original image to produce augmentation-averaged pseudo-labels [36]. However, within Cell Painting research, several standard augmentations have proven detrimental to model performance [30, 15, 8], since altering aspects like image brightness can fundamentally imply that the biological cellular response has also changed. Given the basic flip and rotation augmentations used in our architecture, there is not enough diversity to generate robust teacher pseudo-labels. Test-time template adjuster (T3A) computes a pseudo-prototype representation for each class [14]. Given the limited number of examples per class in both our datasets, as well as the batch effects, these prototypes did not provide robust supports to structure classification around.

### 4.4 Evaluation

#### Perturbation classification accuracy

To characterize model performance, we use average perturbation class classification accuracy – i.e., predicting the genetic perturbation present in each image. We evaluate results using batch-separated splits for both RxRx1 and JUMP-CP where each test batch has not been seen during training. This therefore mimics the scenario whereby new experimental data has been generated for an adjacent task, so a technique like test-time adaptation would allow our previously trained model to adapt to the new batch(es).

Prior research has shown that batch-stratified splits, where the model has seen perturbations from the same batches during training and test time can overstate results and be detrimental to determining how well the architecture overcomes the experimental batch effects which may obscure biological signal [30].

For RxRx1, we use the same training and test splits as the original paper (see 3.1), also repeating each run five times with different seeds to show model performance variance. For JUMP-CP, since there are only five batches, we repeated model training five times, each time using a different batch as the held-out test set. This approach is necessitated by the low number of individual batches in the JUMP-CP data compared with RxRx1.

#### Baseline models

The same baseline models as Sypetkowski et al. [30] are used for the RxRx1 dataset to ensure fair comparability. For JUMP-CP the DenseNet161 architecture specified in Sypetkowski et al. [30] was replicated, confirmed to be accurate based on results on RxRx1, and then that approach was used to set the baseline for the JUMP-CP data. Results significance was assessed via one-sample t-test against the reported AdaBN baseline mean for RxRx1, as individual run results were unavailable from Sypetkowski et al. [30]. However, pairwise comparisons between methods *within* this paper were conducted using Welch’s t-test.

## 5 Results and experiments

### 5.1 Reducing batch effects

#### RxRx1

The results in Table 2 show a consistent improvement in perturbation classification accuracy when employing the architectural and test-time adaptation additions dis-cussed in Sec. 4.1-4.3. Using Tent to minimize entropy loss at inference time while only changing the scale and shift parameters of the model increases test-set accuracy for RxRx1 by 2.0% compared with the prior state-of-the-art AdaBN baseline set by Sypetkowski et al. [30]. When the diversity and pseudo-label classification losses of SHOT are included, the model improves by 2.9% over AdaBN, before our cell count gradient reversal adds an additional 1.6%. Overall, this significantly improved accuracy by 4.5% over the original RxRx1 AdaBN benchmark (t=63.64, p<0.0001). The 1.6% improvement gain from the addition of CCR to SHOT was also assessed as statistically significant (t=9.74, p<0.0001).

**Table 1:** Summary of the two datasets used to test batch effect reduction in this work. Genetic perturbations include knock-down (KD) or knock-out (KO). *The JUMP-CP dataset was subset to only include CRISPR genetic perturbations with ≥ 5 replicates, see 3.2 for more details.

| Dataset | Cell Type | Channels | Perturbation type | # Images | # Batches | # Perturbations |
| --- | --- | --- | --- | --- | --- | --- |
| RxRx1 | U2OS, HUVEC, RPE, HepG2 | 6 | Gene KD (siRNA) | 125,510 | 51 | 1,139 |
| JUMP-CP* | U2OS | 5 | Gene KO (CRISPR) | 21,780 | 5 | 484 |

**Table 2:** Perturbation classification accuracy (%) comparing different approaches to batch correction and test-time adaptation. For RxRx1 the model was trained five times using different seeds and tested on the same test set, for comparability with Sypetkowski et al. [30]. For JUMP-CP the model was tested on each of the five different held-out experimental batches. GR = gradient reversal.

| Method | RxRx1 | JUMP-CP (Subset) |
| --- | --- | --- |
|  | *Perturbation classification accuracy* |  |
| Baseline [30] | 75.1% ±0.2% | 10.4% ±8.3% |
| AdaBN [30] | 87.1% ±0.2% | 28.0% ±7.5% |
| AdaBN + Batch GR [30] | 86.2% ±0.3% | 32.4% ±7.2% |
| Ours (Tent [35]) | 89.1% ±0.2% | 35.0% ±4.9% |
| Ours (SHOT [18]) | 90.0% ±0.3% | 43.4% ±6.4% |
| Ours (SHOT + Batch GR) | 90.1% ±0.2% | 43.2% ±5.9% |
| Ours (SHOT-CCR) | **91.6% ±0.2%** | **43.7% ±5.6%** |

#### JUMP-CP Subset

As expected, the smaller and less diverse JUMP CRISPR dataset was classified with much lower accuracy overall. However, in this scenario, the test-time adaptation methods had a much larger impact on model performance. AdaBN improved classification accuracy by 17.6% over the baseline, while our approach utilizing SHOT and cell count gradient reversal improved classification by a further 15.7%. For this dataset, both gradient reversal methods (batch and cell count) only provide marginal improvement over vanilla SHOT. Figures 3 and 4 contextualize these results. Unlike RxRx1 (Figure 3), the JUMP-CP subset contains only a single cell type across batches with broadly consistent cell count distributions (Figure 4), meaning cell count provides little discriminative batch signal for CCR to suppress. This is consistent with our hypothesis that CCR’s benefit scales with cell count heterogeneity across batches. Crucially, the datasets where this heterogeneity is greatest - multi-cell-type datasets like RxRx1 - are also the most clinically relevant, as generalizing across cell types is a prerequisite for scalable drug discovery pipelines [5, 26].

**Figure 3.**
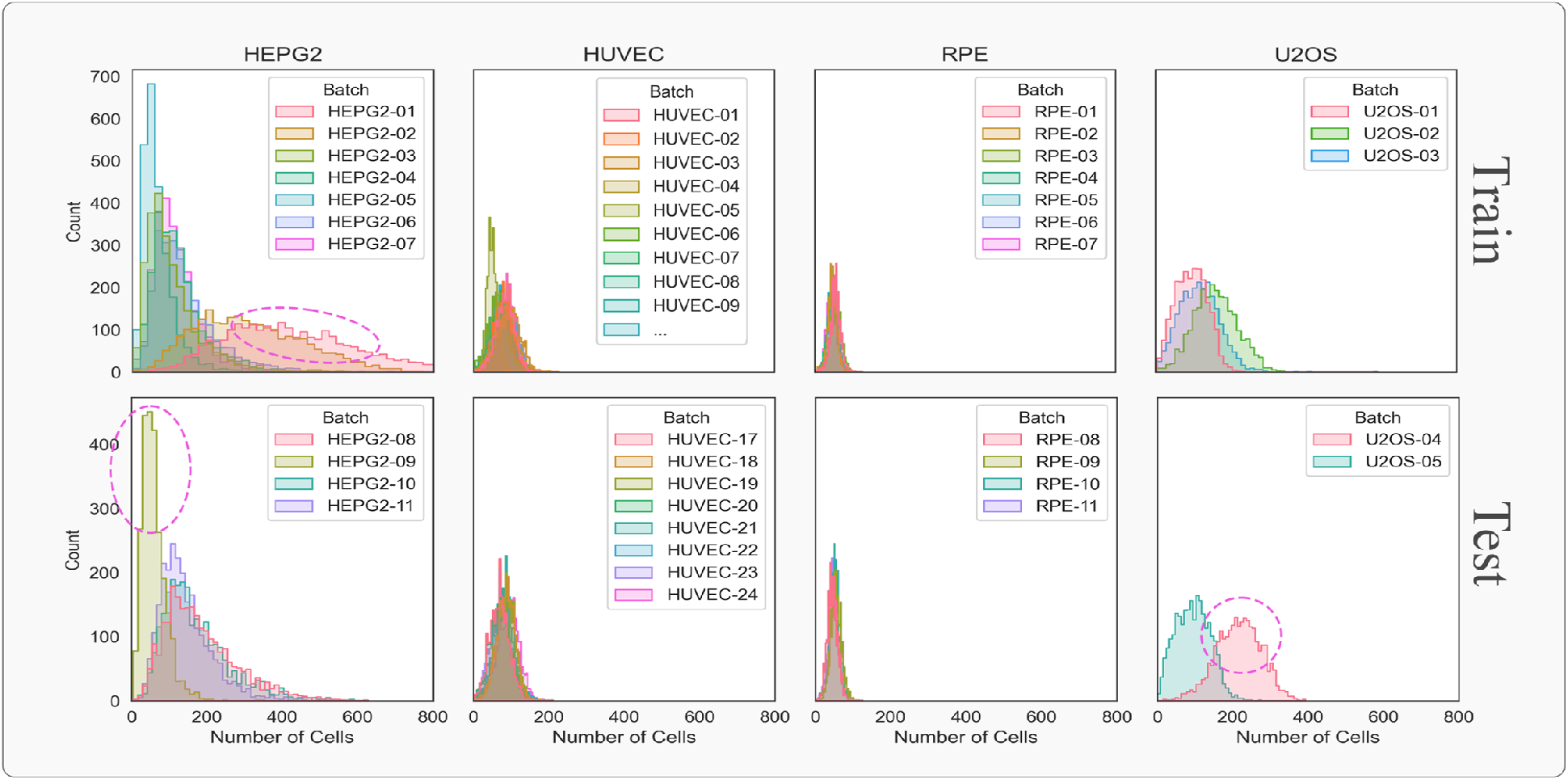
Histograms showing the distribution of the number of cells per field-of-view image within RxRx1. Some batches show clearly different cell count distributions (circled pink) when compared to others of the same cell type. Please zoom in for details.

**Figure 4.**
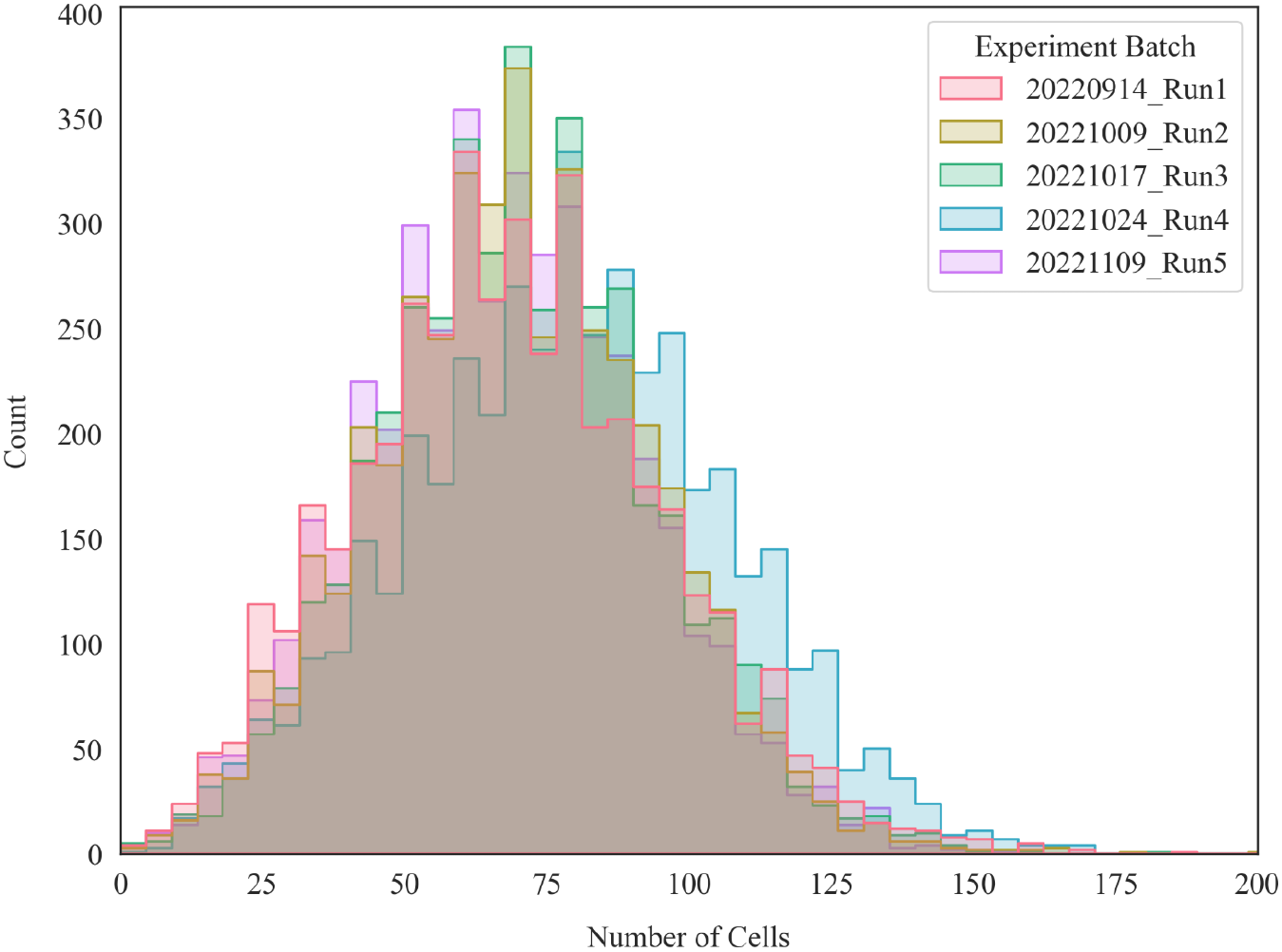
Histograms showing the relatively uniform distribution of number of cells per field-of-view image across the different experimental batches in the JUMP-CP CRISPR subset.

### 5.2 Consistency across cell types

Our results show consistent improvement across all cell types in RxRx1 (Table 3). Encouragingly, performance gain was the strongest in the U2OS cell type (+8.0% accuracy), the cell type with the fewest training batches (3). This was the cell type established as most difficult to predict by Sypetkowski et al. [30]. This result suggests that TTA methods, guided by cell-count gradient reversal, are relatively agnostic to cell type, and can be most impactful where there is limited cell-type-specific training data.

**Table 3:** Perturbation classification accuracy (%) per cell type. Under each cell type we show how many different batches were in the training set compared with the test set, i.e. for HUVEC “16/8” represents 16 training and 8 test batches.

| Method | HUVEC<br>(16/8) | RPE<br>(7/4) | HepG2<br>(7/4) | U2OS<br>(3/2) |
| --- | --- | --- | --- | --- |
| **RxRx1** |  |  |  |  |
| Baseline [30] | 84.2% | 79.0% | 76.2% | 26.1% |
| | $\pm 0.2\%$ | $\pm 0.4\%$ | $\pm 0.1\%$ | $\pm 1.5\%$ |
| AdaBN [30] | 92.1% | 87.2% | 86.2% | 68.2% |
| | $\pm 0.2\%$ | $\pm 0.0\%$ | $\pm 0.2\%$ | $\pm 0.1\%$ |
| Ours (SHOT-CCR) | **95.1%** | **91.9%** | **92.0%** | **76.2%** |
| | $\pm 0.1\%$ | $\pm 0.1\%$ | $\pm 0.2\%$ | $\pm 0.3\%$ |

### 5.3 Gradient reversal

While prior research found gradient reversal to be ineffective when trying to adversarially reduce the batch effect signal by predicting what experimental batch each image was produced in [30], here we show that gradient reversal can be effective if it’s focused on a biological prior – cell count. Consistent with this, CCR showed no significant improvement over vanilla SHOT in JUMP-CP, where cell count distributions are relatively homogeneous across batches (Figure 4), but delivered meaningful gains in RxRx1 where cell count varies substantially across cell types (Figure 3).

Batch effects can encompass many different aspects of the Cell Painting image, including the lab conditions, the technical approaches, and the actual chemical solutions and equipment available [5, 8]. Trying to remove the impact of all these aspects at once in Sypetkowski et al. [30] may have had a detrimental effect on the ability of the embeddings to also represent the biological impact of the perturbation. Similarly, when we set the learning rate too high for our cell count gradient reversal layer, the model learned complete agnosticism towards cell count but no longer functioned at the classification task. Conversely, if the learning rate was too low, the model didn’t strip out enough of the cell count-related batch effect and performance failed to improve.

We found that separate gradient reversal parameters (*α* = 5.0 and regression head learning rate = 10^−5^) were critical for model performance, whereby classification accuracy was dependent on reversal strength. Figure 5 shows how the cell count head regressor MSE loss is impacted by *α*. Other research supports this idea of a trade-off between performance and invariance. Often partial invariance is the optimal solution, as opposed to strong adversarial training which can cause performance degradation [2, 11]. By using separate gradient reversal parameters to control how much cell-count related information is retained by the embeddings, we strike a balance between removing batch-specific noise while preserving task-relevant information.

**Figure 5.**
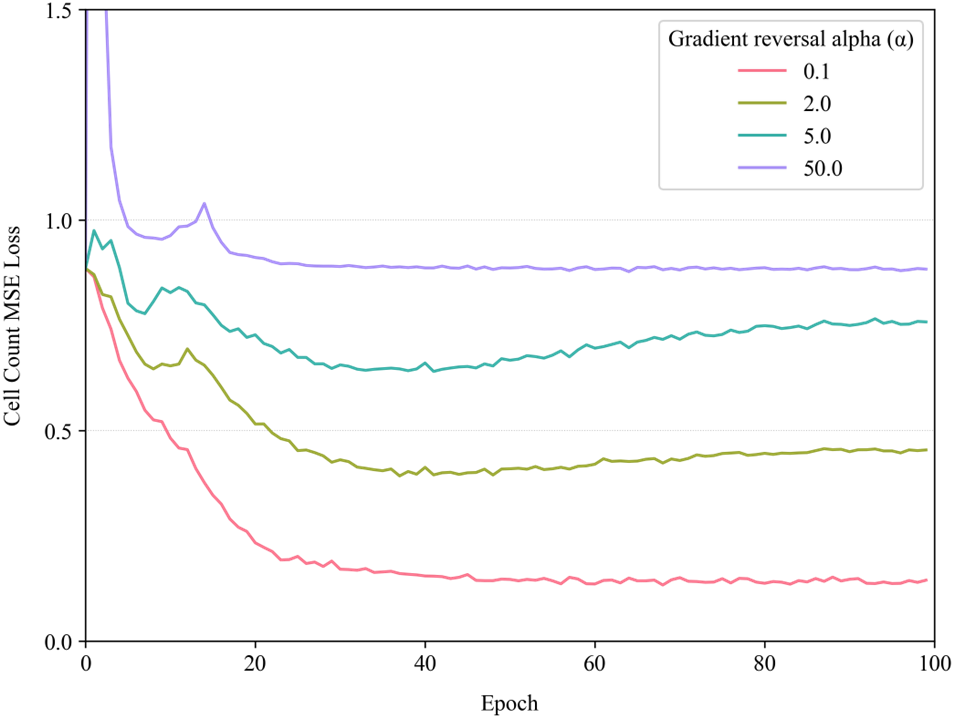
Plot showing the cell count regression head MSE loss throughout training under different gradient reversal *α* parameters. 0.0 loss would indicate the model has learned to perfectly predict the number of cells in each image. Increasing the *α* parameter has a direct impact on how much cell count information is retained in model embeddings.

### 5.4 Model backbone comparison

Table 4 indicates there is very little performance gained from using alternative model backbones. EfficientNetV2 increases AdaBN classification accuracy by 1.7% using 59% more parameters than DenseNet161. ConvNeXt in-creases baseline performance by 1.2% (76.3% compared to 75.1%) while requiring 60 million more parameters than DenseNet161. As expected, since ConvNeXt uses layer (i.e., instance) normalization, AdaBN only impacts the linear layers post the backbone in our setup (see Figure 2) – this causes the performance uplift from AdaBN to be lower with this backbone. These results suggest however that architectural performance improvements are limited and perhaps not worth the additional model size and complexity.

**Table 4:** Perturbation classification accuracy (%) using different pre-trained backbones. # Parameters refers to full architecture.

| Backbone | Method | # Parameters | Pert. accuracy |
| --- | --- | --- | --- |
| **RxRx1** |  |  |  |
| DenseNet161 | Baseline | 41.7m | 75.1% |
| DenseNet161 | AdaBN | 41.7m | 87.1% |
| EfficientNetV2 | Baseline | 66.2m | 71.3% |
| EfficientNetV2 | AdaBN | 66.2m | **88.8%** |
| ConvNeXt (base) | Baseline | 100.0m | 76.3% |
| ConvNeXt (base) | AdaBN | 100.0m | 78.5% |

### 5.5 Cell count and batch selection

Figure 3 shows that there are clear batch effects associated with cell count in RxRx1. We have circled four specific batches across the training and test sets where the cell count distribution is visibly different from other batches performed with the same cell types. Interestingly, the two cell types, HUVEC and RPE, where cell count distributions are almost identical between training and test set are also the best predicted cell types (Table 3).

We observe that for the U2OS cell type, 50% of the test data (Batch U2OS-04) comes from a batch with a different cell count distribution to the other U2OS batches. We investigated the performance of the AdaBN model (chosen to be comparable with Sypetkowski et al. [30]) under three scenarios where all batches with differing cell count distributions, i.e., HEPG2-01, HEPG2-02, HEPG2-09 and U2OS-04, were:

1. All included in the training data only.
2. In the test data only.
3. Excluded entirely from the dataset.

In each case, if one batch was moved from the train or test set, it was swapped with another batch, so the size of the sets remained equivalent (apart from 3. which was smaller due to the excluded batches). By shifting all the batches with markedly different cell count differences to the test set (“AdaBN (shift - test)” in Table 5) we observe improved model performance for the HUVEC and RPE cell types, while the lowest performance is achieved for HepG2 and U2OS.

**Table 5:** Perturbation classification accuracy (%) per cell type when batches with different cell count distributions are shifted either; all to the training set, to the test set, or excluded entirely. Blue: worst performance per cell type. **Black (bold):** best.

| Method | HUVEC | RPE | HepG2 | U2OS |
| --- | --- | --- | --- | --- |
| **RxRx1** |  |  |  |  |
| AdaBN [30] | 92.1% | 87.2% | 86.2% | 68.2% |
| AdaBN (shift - train) | 92.9% | 87.7% | **86.9%** | 75.2% |
| AdaBN (shift - test) | **93.3%** | **88.3%** | 82.2% | 63.6% |
| AdaBN (excluded) | 93.1% | 87.8% | 82.7% | **79.4%** |

When we exclude those batches entirely (“AdaBN (excluded)”), we see the performance for each cell type improve versus the baseline and the performance of U2OS specifically improves by 11.2% to 79.4%. This is despite having two fewer batches in the training data (HEPG2-01, HEPG2-02).

These results further support that cell count is an integral feature to consider when selecting train/test splits, and that including batches with significantly different distributions compared to others of the same cell type may lead to a less robust classifier. Whilst this diversity may mimic real life, the distribution disparity would likely warrant further investigation of the batches in question to ensure there is no technical error that has led to this difference. Our results also suggest that U2OS is not necessarily a more difficult cell type to predict as hypothesized in Sypetkowski et al. [30], more the results are both a function of a lack of data combined with differing test set distribution. This is further supported by comparing perturbation classification accuracy by batch between AdaBN and SHOT-CCR (Figure 6). SHOT-CCR shows the greatest improvement in the *test* batches identified in Figure 4 with differing cell count distributions to the *training* data (HepG2-09, U2OS-04 and U2OS-05).

**Figure 6.**
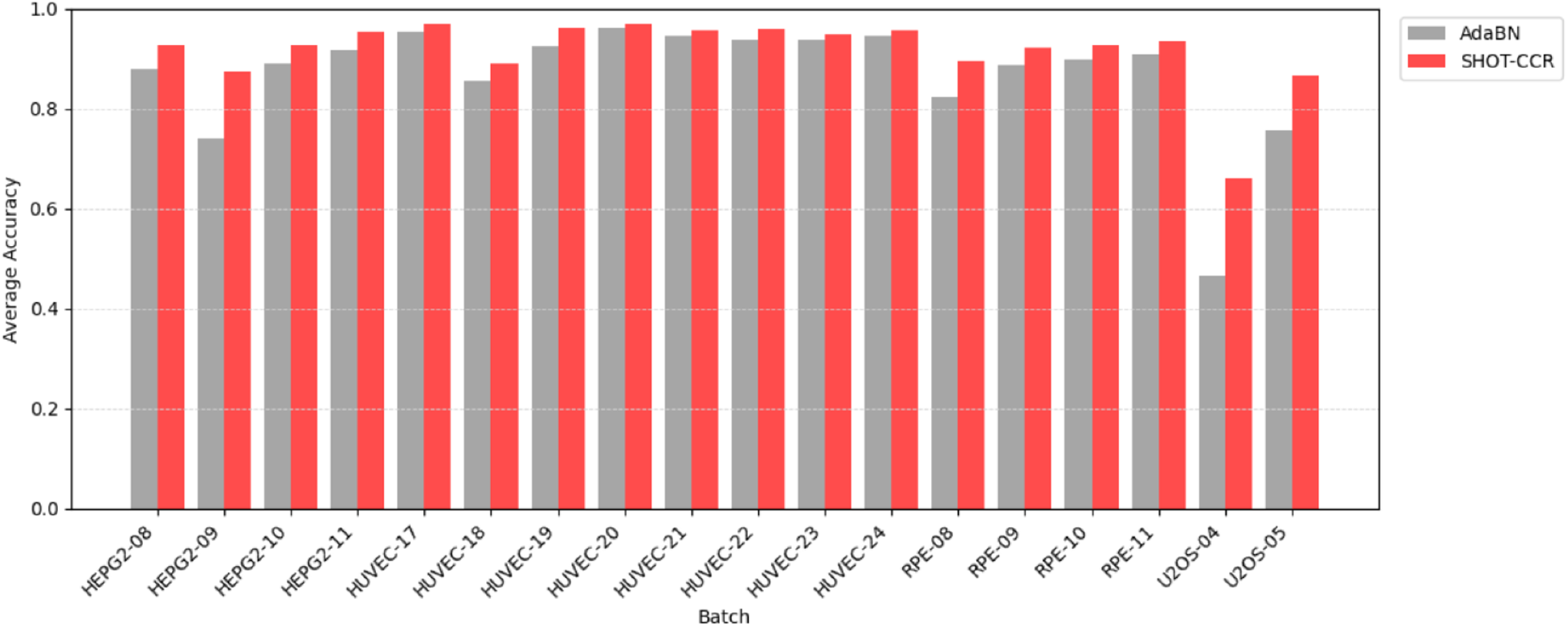
Bar chart showing average perturbation accuracy per test batch, comparing AdaBN with SHOT-CCR. Accuracy improves the most in batches HepG2-09, U2OS-04 and U2OS-05, where data is either distributed differently between the training and test data, and/or is scarce.

### 5.6 Model ablations

Table 6 presents an ablation study across both multi-domain and single-domain (BEN [20]) training and inference settings. The results first highlight that BEN sampling and AdaBN are interdependent — while single-domain sampling alone slightly reduces baseline performance (69.4% vs 75.1%), it is a prerequisite for AdaBN to function correctly, and the combination improves strongly to 87.1%. Notably, general batch-identity gradient reversal (Batch GR) proves actively harmful in the multi-domain setting, reducing accuracy by 3.9% from the baseline, directly supporting the argument that indiscriminate batch-effect reversal strips out task-relevant signal alongside noise. In contrast, CCR alone provides a modest but positive standalone contribution (+1.5% over the single-domain baseline), demonstrating that biologically targeted gradient reversal has value independent of the TTA pipeline. The complementarity of CCR is further evidenced by the small but consistent gain when combined with AdaBN (87.3% vs 87.1%), suggesting CCR is not specific to SHOT. Comparing SHOT-CCR against SHOT + Batch GR (91.6% vs 90.1%) demonstrates that biological targeting of a specific confounder outperforms generic batch gradient reversal by 1.5%. Overall, the table shows incremental progression from 69.4% to 91.6%, with each component contributing positively, validating the design choices of the full SHOT-CCR framework.

**Table 6:** Perturbation classification accuracy (%) on RxRx1 comparing ablations across multi-domain and single-domain (BEN [20]) training/inference batch settings.

| Method | Perturbation classification accuracy |
| --- | --- |
| **RxRx1** |  |
| *Multi-domain training/inference batches* |  |
| Baseline [30] | 75.1% |
| + Batch GR | 71.2% |
| + CCR (Ours) | 73.6% |
| + SHOT [18] | 87.0% |
| + SHOT-CCR (Ours) | 86.9% |
| *Single-domain training/inference batches (BEN [20])* |  |
| Baseline [30] | 69.4% |
| + CCR | 70.9% |
| + AdaBN [30] | 87.1% |
| + AdaBN + Batch GR [30] | 86.2% |
| + AdaBN + CCR | 87.3% |
| + SHOT [18] | 90.0% |
| + SHOT [18] + Batch GR | 90.1% |
| + SHOT-CCR (Ours) | **91.6%** |

### 5.7 Gene enrichment

To assess whether our approach recovers meaningful biological signal, we performed gene set enrichment analysis. We applied g:Profiler [25] to the RxRx1 knocked-down genes where SHOT-CCR showed at least 10% classification accuracy improvement over AdaBN across replicates. Gene enrichment analysis tests whether sets of genes are statistically overrepresented in specific biological pathways or cellular functions compared to random expectation. By applying this test to genes where classification performance has improved specifically, we can determine whether performance gains are concentrated in functionally related categories, which would suggest an improved ability to recover biologically meaningful signal.

We found significant enrichment across multiple categories: cytoplasm (GO:0005737, *p* = 1.5 × 10^−9^), cytosol (GO:0005829, *p* = 2.8 × 10^−3^), helicase activity (GO:0004386, *p* = 1.1 × 10^−2^), nucleic acid conformation isomerase activity (GO:0090729, *p* = 1.1 × 10^−2^), and endomembrane system (GO:0012505, *p* = 1.3 × 10^−2^). The genes include multiple DEAD-box RNA helicases, vesicle trafficking components, ER-associated proteins, and endosomal sorting factors.

The enrichment in RNA helicases and nucleic acid processing machinery is particularly informative. These genes affect nucleolar morphology and cellular stress responses through mechanisms that produce morphologically subtle phenotypes compared with more dramatic perturbations like complete cytoskeletal disruption [31]. These subtle phenotypes are particularly susceptible to being obscured by batch effects.

We also observe strong enrichment for endomembrane system components and cytoplasmic localization, cellular compartments explicitly targeted by dedicated Cell Painting channels. While cytoplasmic GO terms encompass thousands of genes, the concurrent enrichment of more specific categories like endomembrane system and helicase activity supports genuine underlying biological patterns being recovered rather than an artifact of pathway size. This suggests that our method improves classification not only for morphologically subtle phenotypes, but also for prominent cellular structures where batch-to-batch technical variation can corrupt morphological readouts.

## 6 Conclusion and future directions

In this paper we have demonstrated that adversarial training targeting a biologically motivated confounder, cell count, combined with test-time adaptation methods, can effectively dampen batch effects in Cell Painting data and improve perturbation classification accuracy across multiple cell types and datasets, outperforming the previous benchmark of Sypetkowski et al. [30]. We set out the framework by which this approach can be applied to other Cell Painting datasets well as providing advice regarding dataset composition, specifically how differing cell count distributions across experimental batches can impact both model training and inference. Where there are substantially different distributions of a biological indicator such as cell count across batches containing the same cell type, we would encourage researchers to consider this in both their approach and in the discussion of results.

Lastly, we have successfully transferred our approach to a separate, major, Cell Painting dataset (JUMP-CP), where test-time adaption methods again proved to be significantly beneficial for classification accuracy. We sub-selected a suitable cohort of samples from JUMP-CP which are comparable with RxRx1 as it contains 484 overlapping gene targets (although modified with CRISPR knockout rather than siRNA knockdown). The JUMP-CP subset used in this work can be used as an additional benchmark in the field going forward.

### 6.1 Future directions

We would encourage future work to investigate whether the test-time adaptation techniques which proved successful in this work can be adapted to transformer-based models by altering layer normalization rather than batch normalization layers.

Furthermore, cell count is only one aspect of batch effect obfuscation, there are many more which may ultimately benefit the model if their affects are reduced [1, 8, 26]. For example, Yan et al. [37] show triple batch effects in CellProfiler Cell Painting data, being the experimental batch, as well as the column and row of the plate the experiment was performed on. It may be beneficial to use separate parameters to control the strength of the gradient reversal for these different types of confounding factors.

Finally, we would recommend researchers keep adding results from more large-scale Cell Painting datasets so that we eventually have a comprehensive environment to rigorously test all conceived methods against. This would enable us to collectively move towards a foundational method for dealing with microscopy batch effects.

## Acknowledgments

This work was supported by the UKRI/BBSRC Collaborative Training Partnership in AI for Drug Discovery, led by Recursion Pharmaceuticals in partnership with Queen Mary University of London. We are extremely grateful to Maciej Sypetkowski for his guidance and help throughout this project, especially when replicating the original results from RxRx1. The authors would like to thank the Broad Institute and the JUMP Consortium for making the JUMP-CP Cell Painting data freely available and accessible. We would also like to thank Safiye Celik and Inês Sequeira for their valuable feedback on the draft manuscript.

This work was supported by the Engineering and Physical Sciences Research Council [grant number EP/Y009800/1], through funding from Responsible Ai UK (KP0016). This work acknowledges the support of the National Institute for Health Research Barts Biomedical Research Centre (NIHR203330).

This research utilized Queen Mary’s Apocrita and Andrena HPC facilities, supported by QMUL Research-IT. http://doi.org/10.5281/zenodo.438045.

